# AlcoR: alignment-free simulation, mapping, and visualization of low-complexity regions in biological data

**DOI:** 10.1101/2023.04.17.537157

**Authors:** Jorge M. Silva, Weihong Qi, Armando J. Pinho, Diogo Pratas

## Abstract

Low-complexity data analysis is the area that addresses the search and quantification of regions in sequences of elements that contain low-complexity or repetitive elements. For example, these can be tandem repeats, inverted repeats, homopolymer tails, GC biased regions, similar genes, and hairpins, among many others. Identifying these regions is crucial because of their association with regulatory and structural characteristics. Moreover, their identification provides positional and quantity information where standard assembly methodologies face significant difficulties because of substantial higher depth coverage (mountains), ambiguous read mapping, or where sequencing or reconstruction defects may occur. However, the capability to distinguish low-complexity regions (LCRs) in genomic and proteomic sequences is a challenge that depends on the model’s ability to find them automatically. Low-complexity patterns can be implicit through specific or combined sources, such as algorithmic or probabilistic, and recurring to different spatial distances, namely local, medium, or distant associations.This paper addresses the challenge of automatically modeling and distinguishing LCRs, providing a new method and tool (AlcoR) for efficient and accurate segmentation and visualization of these regions in genomic and proteomic sequences. The method enables the use of models with different memories, providing the ability to distinguish local from distant low-complexity patterns. The method is reference- and alignment-free, providing additional methodologies for testing, including a highly-flexible simulation method for generating biological sequences (DNA or protein) with different complexity levels, sequence masking, and a visualization tool for automatic computation of the LCR maps into an ideogram style. We provide illustrative demonstrations using synthetic, nearly synthetic, and natural sequences showing the high efficiency and accuracy of AlcoR. As large-scale results, we use AlcoR to unprecedentedly provide a whole-chromosome low-complexity map of a recent complete human genome and the haplotype-resolved chromosome pairs of a heterozygous diploid African cassava cultivar.The AlcoR method provides the ability of fast sequence characterization through data complexity analysis, ideally for scenarios entangling the presence of new or unknown sequences. AlcoR is implemented in C language using multi-threading to increase the computational speed, is flexible for multiple applications, and does not contain external dependencies. The tool accepts any sequence in FASTA format. The source code is freely provided at https://github.com/cobilab/alcor.

## Introduction

With the current unprecedented developments in sequencing techniques and reconstruction methodologies, the availability of complete sequences with high-quality and without gaps is steadily increasing [Miga et al., 2020, Wang et al., 2022, Qi et al., 2022], providing direct applicability to the accurate analysis of these sequences, specifically large-scale data complexity studies. However, the current methodologies are not fully prepared to deal with the scale and characteristics of the reconstructed data, becoming prohibitive regarding computational resources and without the desirable precision and accuracy.

Several considerations must be inquired to address the problem adequately; for example, consider the first genomic or proteomic sequence that comes to mind. Does a method to accurately distinguish low-complexity regions (LCRs) from the remaining sequence exist? Does this method provide the capability to address different characteristics and proximity within these patterns? If all yes, can this method be implemented as an efficient tool? Can it be implemented to provide visualization of the localized regions? How can this method be appropriately benchmarked? These are the five main questions raised and addressed in this article.

We refer to LCRs as the sub-sequences from a sequence of symbols that are considered to contain lower complexity or a higher number of repetitive characteristics measured above a certain threshold. As an example, consider the following sequence: “ATGCTCGAAAAAAAAAACGAGCAT”. From this sequence, is there any repetitive or low-complexity region? One can immediately identify the low-complexity region as the repetition of “AAAAAAAAAA” and the corresponding region coordinates of 8 to 17. However, although not immediately recognizable, there are other LCRs. Specifically, the “ATGCTCG” sub-sequence from positions 1 to 7 is an inversion of the “CGAGCAT” sub-sequence at positions 18 to 24. Inversions are sub-sequences from an original one where instead of a normal order are in reverse order (from the end to the beginning or the opposite), followed by a substitution where each element becomes exactly another one (A becomes T, C becomes G, G becomes C, and T becomes A). Despite the simplicity of this transformation, if the method that performs this type of detection is not prepared to find this algorithmic characteristic, the low-complexity region will not be classified as such. Therefore, the better the model, the higher the accuracy and the cumulative amplitude of these regions.

Many other types of low-complexity patterns may be contained in sequences, for example, tandem repeats, exact or similar genes, poly-tales, GC regions, and hairpins, among others [Hubbard et al., 2002]. However, low-complexity sequences can cause serious challenges for search and clustering algorithms, for example, based on matching words or patterns. Detecting these regions in biological sequences is also related to differences in the accuracy of sequencing technologies [Reis et al., 2022] and assembly strategies [Qi et al., 2022] to reconstruct the original biological sequence. In fact, there are many highly important applications that depend on the correct and non-underestimation of these regions. For example, the recent state-of-the-art viral integration caller uses LCRs as part of fundamental analysis in integration classification [Rajaby et al., 2021]. Specifically, inaccuracy on previous integration callers is mainly caused by the incorrect alignment of reads in repetitive regions, leading to false positives. This classification is of extreme importance because predicting virus integration can help uncover the mechanisms that lead to many devastating diseases [Kao and Chen, 2002, Schiffman et al., 2007, Parkin, 2006, Xu et al., 2019].

However, building a model that can accurately identify all these LCRs is not a trivial process because, generically, biological sequences contain LCRs that may be characterized by mutations, heterogeneity, and non-stationary content, and additionally, noise and errors that may be inserted through the sequencing and reconstruction (assembly) process, analogous to predict outcomes based on imperfect data [Golan, 2018]. Moreover, the size of the sequence also matters. If we consider now that there were twenty million random symbols between the identified inverted regions of the above example, would the model still have enough memory and precision to identify it as a low-complexity region? What about the number of sequences that would have exact or similar sequences? There is a high probability of having more 7-nucleotide sequences exactly as “ATGCTCG” or “CGAGCAT” (assuming a uniform distribution) for approximately twenty million symbols. Therefore, is it considered an underestimation if the model considers these sub-sequences of higher redundancy? Probably, yes. So, where to draw the line of underestimation? Fortunately, models that are upper-bounded approximations to the Kolmogorov complexity implicitly respect this property [Li and Vitányi, 2008].

Specifically, the Kolmogorov complexity is defined as the size of a shortest program that represents a particular sequence and halts [Kolmogorov, 1965]. Therefore, the Kolmogorov complexity is a shortest quantity measure but non-computable that can only be upper-bounded by an efficient model. Such a model can be a lossless data compressor because its main aim is to minimize the data representation without loss of information. However, the quality and efficiency of the compressor dramatically influence the accuracy and precision of the predictions. Therefore, data compressors that are prepared to deal with specific properties of the data and consistently provide higher compression ratios are better candidates for predicting LCRs through the generation of complexity profiles.

In practice, a complexity profile is a numeric sequence that estimates the amount of information (for example, in bits) to compress or represent each symbol of the sequence. The size of the complexity profile is equal to the size of the sequence and, through vertical alignment, provides immediate localization of complexity differences between the sequence regions found by the data compressor in use [Pinho et al., 2013]. Regions below a certain threshold are considered low-complexity or to contain higher redundancy.

The literature provides several examples of the computation of complexity (or entropy) profiles or equivalent approaches for many applications [Vinga and Almeida, 2007, Kempa and Prezza, 2018]. For example, in the area of computational security [S Resende et al., 2019], these profiles have been used for mimicking anti-viruses [Menéndez and Llorente, 2019] and detecting malware [Alshahwan et al., 2020]. Although the success in the application of other distinct areas, namely with the classification of galaxy clusters [Donahue et al., 2006] and finding characteristic fingerprints in lithium-ion cells [Osswald et al., 2015], the use of these profiles has been widely applied to biological sequence analysis [Allison et al., 1998, Rivals et al., 1997, Crochemore and Vérin, 1999]. Examples of these biological applications are regulatory regions prediction [Allison et al., 2000, Dix et al., 2006, Pinho et al., 2013, Wu et al., 2019], identification of inversions [Pinho et al., 2011a, Hosseini et al., 2017], predicting coding regions [Pinho et al., 2006], diversity estimation [Chao and Jost, 2015], prediction of mRNA and lncRNAs folding [Lai et al., 2018, Ermolenko and Mathews, 2021], rearrangements detection [Hosseini et al., 2020, Jiang et al., 2020], improving genome reconstruction [Pratas et al., 2020], and to predict local LCRs [Troyanskaya et al., 2002].

The method proposed in this article (AlcoR), with description and formalization provided in the next section, uses a bidirectional compression scheme of an input string assuming two causal directions, from the sequence’s beginning to end and the opposite, followed by its minimum, smoothing, segmentation, and visualization operations. The method uses a compression scheme that combines multiple context models with specific memory capacities to consider different distances between patterns. Moreover, a free and efficient standalone implementation of the method is provided in C language with the additional flexibility to address challenges in other areas.

To benchmark the method, we use synthetic sequences as one of the most effective practices in benchmark-test methods and data analysis [Escalona et al., 2016, Posada, 2020]. Here, synthetic sequences are defined as computer-generated sequences that resemble one or multiple features of natural sequences. Synthetic sequences provide a controlled environment with complete or partial expected outcomes, similar to a gold standard. Additionally, they can be used to create or identify protein sequences with optimized properties [Huang et al., 2016, Yang et al., 2019, Wu et al., 2021]. Ideally, they should be used as the first benchmark test for most existing computational biological tools, especially in low-complexity analysis, given structural complexity differences.

The literature contains several methodologies to generate synthetic sequences both using models with low to high simplicity for genomic and proteomic sequence [Wu et al., 2021, Feldkamp et al., 2001, Ponty et al., 2006, Rouchka and Hardin, 2007, Chen et al., 2009, Pratas et al., 2011, Angermueller et al., 2019, Rong et al., 2021]. This generation has been augmented to specific file formats that additionally simulate the process of NGS methodologies (e.g., [Huang et al., 2012]), providing a critical application to compare existing and new NGS tools or analytical pipelines [Escalona et al., 2018].

Specifically, synthetic sequences can contain different characteristics and generation sources. For example, a synthetic sequence can be generated according to a specific distribution, assuming statistical independence between the outcome of the symbols, or generated according to a model that has been trained in one or multiple *natural* DNA sequences. These sequences can also contain deliberated rearrangements (e.g., translocation, inversions, fusions) or a small variation, substitutions, additions, or deletions of one or multiple bases. Another feature that synthetic sequences can contain is the heterogeneity created by multiple sources, for example, through the alternated combination of segments produced by a simple independent model and a model trained with a particular sequence. As an analogy, it can be seen as a book independently written by different authors and combined using a conjunction of pages.

Therefore, to benchmark the method for locating LCRs, we also provide a flexible method that enables the simulation of synthetic sequences. In the next section, we describe this methodology along with the details of its implementation into an efficient tool. Moreover, several illustrative demonstrations are provided using synthetic, nearly-synthetic, and natural sequences with different characteristics. Although the AlcoR method is able to provide additional statistical and algorithmic analysis than common alignment mapper tools, we provide several illustrative comparisons for validation and benchmark purposes.

After the benchmark, we apply the LCRs mapper to large-scale biological data, providing the complete low-complexity maps for a complete human genome and a complete diploid cassava genome, including a comparison with RepeatModeler and RepeatMasker [Flynn et al., 2020, Smit, AFA and Hubley, R and Green, P]. Using both synthetic and natural sequences, we provide evidence that the AlcoR tool can efficiently and accurately model probabilistic and several algorithmic patterns, for example, with a considerable degree of substitutions and inversions, unveiling several structural characteristics of sequences without prior knowledge or references.

## 1 Methods

This section describes the methods used to map, extract, and visualize LCRs in biological sequences. Additionally, we develop and provide a method in this section to simulate synthetic or nearly-synthetic sequences for the main method benchmark purpose. Finally, we provide the respective implementation details of the whole set of tools. The details of the parameters and how to reproduce the methods are available in Supplementary Sections 2 and 3.

### 1.1 LCRs mapper

The proposed method in this article uses lossless data compression for estimating the complexity profiles for further segmentation of LCRs. The biological sequences data compression field has now been three decades and started with Biocompress [Grumbach and Tahi, 1993]. Afterwards, many algorithms emerged, mostly modelling the existence of exact or approximate repeated and inverted repeated regions, through the usage of simple bit encoding, dictionary approaches, or context modelling; a few examples are [Grumbach and Tahi, 1994, Manzini and Rastero, 2004, Cherniavsky and Ladner, 2004, Korodi and Tabus, 2005, Cao et al., 2007, Mishra et al., 2010, Rajeswari and Apparao, 2010, Gupta and Agarwal, 2011, Pinho et al., 2011b,c, Pratas et al., 2016, Kryukov et al., 2019, Liu et al., 2020, Grabowski and Kowalski, 2022, Deorowicz et al., 2023].

Although any efficient lossless data compression method that can provide the number of bits to compress a given symbol is a candidate, a compression scheme that proved to provide state-of-the-art results in genomic sequences in balance with affordable computational resources trade-off is used [Silva et al., 2020, 2021, Kryukov et al., 2020]. This compression method uses soft-blending between context models (FCMs) of several context depths and substitution context models with specific sub-programs, in the case of genomic sequences, to handle inversions [Pratas et al., 2019].

The compression method already provides a setup with several parameterized models for non-expert usage. Nevertheless, it permits to customize the number of models, context depths, mixture approach, and memory capacity, among others.

Figure 1 illustrates the main steps in this methodology. Accordingly, the complexity profiles are generated using a bidirectional compression scheme of an input sequence assuming two causal directions, from the sequence’s beginning to the end (left-right) and the opposite (right-left), followed by the minimum of both directions, smoothing, and segmentation of the regions below a certain threshold. Finally, the visualization of the regions through the generation of a map automatically occurs. Although we use sequence simulation (generation) to benchmark this methodology, the simulation and the extraction of the sequences are optional operations.

**Figure 1.**
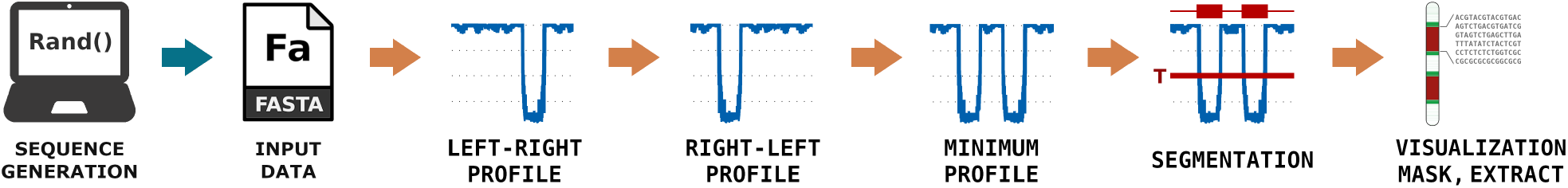
AlcoR pipeline for mapping and visualization of LCRs of a sequence using the bidirectional complexity profiles followed by the minimum, segmentation, and automatic low-complexity map generation (visualization; SVG format). The mask and extract operations of the LCRs and the simulation of synthetic sequences are optional.

The bidirectional computation is key in this process, ensuring that even an *original* sub-sequence that is a copy of another is efficiently identified as a LCR. If the previous section’s exemplifying sequence was used to generate the complexity profile assuming only the beginning to the end order, the first seven symbols would not be identified as a low-complexity region. This feature is a characteristic of models that do not track the positions where copies occur, mainly to save computation time and memory. Therefore, the two directions are processed in parallel to reduce the computational resources substantially.

#### 1.1.1 Formalization and parameters

Consider a source *X*, that has generated *n* symbols from a finite alphabet Θ with size |Θ|. The nature of the source, *X*, is unknown but the sequence of symbols *x*_0_, *x*_1_, …, *x*_*n−*1_ is known. The computation 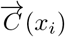 delimits the compression of left-to-right symbol access, i.e. from *x*_0_ to *x*_*n−*1_ by numeric increasing order, and 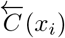delimits the compression of right-to-left access, i.e. from *x*_*n−*1_ to *x*_0_ by numeric decreasing order.

The employed data compressor, *C*(*x*), is characterized by a combination of multiple finite-context models [Pinho et al., 2010, Pratas et al., 2016] (FCM). An FCM has the Markov property, in which the conditional probability distribution of observing a symbol, *R*, depends only on the state of the preceding *k*-mer, respecting the following equality

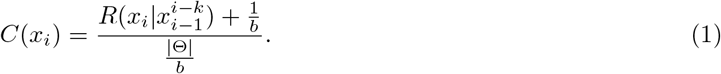

Notice that *b ≥* 1 corresponds to a value that is associated with a bet; where the higher the number, the higher the probabilistic confidence (higher bet). Therefore, when *b* = 1 turns into the Laplace estimator, otherwise behaves progressively as a likelihood estimator.

The FCM models can be combined with other models, such as extended finite-context models [Carvalho et al., 2018] or the substitution tolerant context models [Pratas et al., 2017]. The combination of these models can only access specific memory capacities and, hence, consider different distances between patterns. The access to the sequence is given by

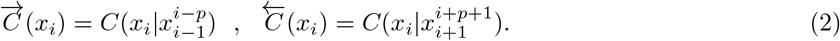

Accordingly, the memory used to independently compute 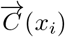 and 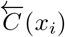 is characterized by a cache history of *p* symbols. This means that the models do not consider older outcomes from the sequence that are far from *p* symbols. Accordingly, at symbol position *i* the model of 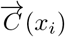 can predict the information content having access exclusively to 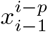 as *x*_*i−*1_, …, *x*_*i−p*_, where for *i < p* the lowest *i* = 0. For predicting the 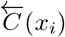 the model has exclusively the access to 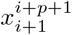 as *x*_*i*+1_, …, *x*_*i*+*p*_, where for *i > p* then *i* = *n −* 1. Both computations run in parallel because the models are independent and it improves substantially the real processing time.

Naturally, the minimum of the two profile computations creates a small degree of underestimation. Despite this degree of underestimation, the impact in the large majority of the cases is negligible for typical thresholds. Additionally, it can be considered affordable because the main idea is to identify the low complexities independently if a region is the original proceeded by a copy.

To compute the minimal bi-directional complexity profiles, both left-to-right as 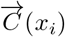 and right-to-left as 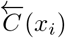, computations are applied according to

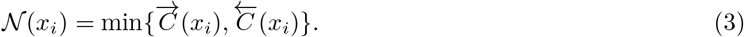

Specifically, both 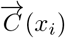 and 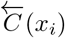 computations run in a different thread, and then the minimum of both is computed when both threads end the computation. After, the *𝒩* (*x*_*i*_) output is averaged according to the following moving average

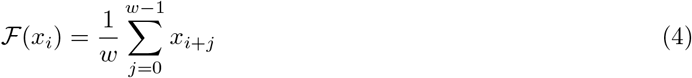

where *w* is the window size parameter *∈* IN^+^.

The initial and final positions of each region below the threshold, *𝒯*, are provided as standard output. By default, the threshold is

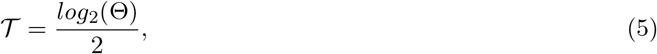

but *T* can also be set to a custom value, where *𝒯 ∈* IR^+^.

Finally, the method outputs the LCRs below the threshold, where regions shorter than a particular size (*s*) can be automatically discarded from the final set.

### 1.2 Extraction, Information, Masking, and Visualization of LCRs

The AlcoR methodology enables to extract LCRs of the input FASTA file using the coordinates generated from the mapper. This characteristic allows further analysis using other approaches, namely comparing or classifying the sequence content within multiple regions.

Another feature that AlcoR provides is to compute the length (nucleotide or amino acid sequence) and the percentage of G and C bases that are registered along with the coordinates of each FASTA read. This is a fast and simple implementation that allows extended analysis for comparative purposes. For example, to understand the trends of LCRs.

A very useful feature of AlcoR is the ability to soft mask the sequences contained in the FASTA file, where each sequence corresponding to a LCR previously mapped is transformed into lowercase symbols.

Moreover, AlcoR enables the automatic generation of an image file (SVG format) depicting the LCRs in scale with the amplest sequence using an ideogram style. We call this functionality the visualization; it enables reading multiple LCRs from multiple sequence files, additionally providing the capability to combine different levels of low-complexity in a single image through the concatenation of the output mapped regions. Additionally, the visualization method enables setting the thickness and distance between the bars and the color of the regions, background, and border. Strict or round corners are also an option. Finally, it provides the functionality to synthetically enlarge the regions to increase visibility when the number of regions is low, or the length is small.

### 1.3 Simulation of sequences

The simulation of synthetic sequences provides a controlled environment to benchmark-test the AlcoR mapper or any other tool that uses FASTA data. Accordingly, we developed a FASTA simulation tool to deal with three main types of sub-models, namely file extraction, pseudo-random generation, and modeling generation. The file extraction simulation mode considers a sub-sequence from a FASTA file containing a sequence using the initial and ending positions. The pseudo-random generation simulates custom sequences using a Linear congruential generator (LCG) with seed and size as the main parameters. The modeling generation, learns from a FASTA file and further generates a sequence with a custom size using a finite-context model of a given context order and bet parameter. This model will store the counters seen in a particular sequence (or set of sequences) and, then, it will compute equation 1, but instead, it will generate a *y* using 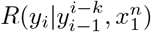. For each simulation model a specific FASTA file and FCM are used, providing flexibility to include modelling using different FASTA files, context orders, and parameters.

In any of the sub-sequences simulated by the previous models, specific transformations can be applied, namely inversions and mutations. The inversion considers the input sequence is reversed and, in the case of DNA, complemented. The mutations applies SNP (Single Nucleotide Polymorphism) mutations using specific intensities, according to a probability of specific mutation, through the generation of pseudo-random symbols (LCG). The SNP mutations can be independently applied for substitution, deletion and addition types.

A very important feature of this method is the capability to generate multiple sequences with different characteristics in a single FASTA file. Therefore, the content of the file can have multiple combinations of the previously reported functionalities through the conjunction (automatic concatenation of the content), enabling it to provide extremely high flexibility.

Additionally, this method enables the application of protein sequences and other symbolic sequences as long as they respect the FASTA file format. Moreover, the alphabet of the sequences can be selected or reduced to ignore certain symbols and provide custom synthetic sequences.

### 1.4 Implementation

The AlcoR method is implemented in C language and does not contain any external dependencies, including the SVG map generation. The source code and benchmark scripts are freely provided at the repository https://github.com/cobilab/alcor. Additionally, we provide a website with several pipelines, including a video to better understand AlcoR, available at https://cobilab.github.io/alcor.

The AlcoR tool contains one main menu (command: AlcoR) with five sub-menus for computing the features that it provides, namely info, extract, mapper, simulation, and visual. The info tool retrieves information of the length and GC percentage for each FASTA read. The extract tool extracts a sequence of a FASTA file using positional coordinates (independent from the existing headers of the FASTA files). The mapper tool computes the LCRs of a FASTA read while providing bidirectional complexity profiles and further structural similarity analysis - this option includes soft masking the LCRs of a FASTA file. The simulation tools provides FASTA sequence simulation with features: file extraction, random generation, sequence modelling (with SNPs specific mutations). Finally, the visual tool computes an SVG file with the respective map containing the LCRs.

For each of the sub-menu tools, supplementary section 3 provides details on each feature, with special focus to the most important parameters.

## 2 Results

Benchmarking methods for unsupervised identification of LCRs is challenging because comparing existing methods requires equivalent characteristics and objectives. Specifically, the number of existing methods for biological sequences is low, and only a few provide computational tools that can be effectively installed and run. Some examples of these tools for DNA sequences are Cafefilter [Williams, 1998], DUST [Morgulis et al., 2006] while for protein sequences SEG [Wootton and Federhen, 1993] and CARD [Shin and Kim, 2005]. For specific purposes, there are several tools, for example, RepeatModeler for the automated genomic discovery of transposable element families [Flynn et al., 2020] and RepeatMasker for masking LCRs [Smit, AFA and Hubley, R and Green, P].

From this set, some are not prepared to deal with FASTA data, the computational resources of the methods needed to handle these sequences are prohibitive, or the current data size is not appropriate for the computation or visualization. Also, since some of them have been built for specific uses, their comparison becomes highly subjective. Moreover, the AlcoR method contains a low-complexity analysis that comprises local and distant regions at the same time, currently providing singular characteristics. Therefore, to fairly benchmark AlcoR, we progressively provide several demonstration tests using synthetic, nearly synthetic, and natural sequences. In the latter, we provide comparative results with the NCBI sequence viewer (NCBIv) [Rangwala et al., 2021] which is one of the most compelling and used online tools. Finally, we provide the complete maps of the recent complete human genome and the diploid Cassava chromosomes, including a comparison with RepeatModeler and RepeatMasker for the Cassava. These results can be reproducible using the repository provided at https://github.com/cobilab/alcor.

All the computations have been carried out using a laptop computer with 8GB RAM, eight 11th Gen Intel Core i5 @ 2.40GHz, an SSD disk with a capacity of 512 GB, and running as an operating system the Linux Ubuntu 20.04 LTS.

### 2.1 Demonstrations

This subsection provides a set of illustrative and interactive demonstrations of the method using several synthetic, nearly synthetic, and natural sequences characterized by different types of copies and transformations, namely inversions and mutations. These demonstrations provide the ability to change conditions to provide personalized analysis anytime during these results, for example, the edition of the smoothing window size, threshold, compression models, or the simulation composition. The conjoint computation of the whole demonstrations described below took merely a few seconds using the computational resources described above. Additional information on the instructions to run the results is available in Supplementary Section 2, while detailed information on the AlcoR parameters and meaning is available in Supplementary Section 3.

#### 2.1.1 Demonstration A

The first demonstration (demo) contains two scripts (GenSeqA.sh and DemoA.sh) that can be accessed using the folder “demos/demoA/”. This demo uses AlcoR to simulate a FASTA file (sampleA.fasta) through the recursive generation of several pseudo-random (synthetic) sequences. For generating this file, the command “./GenSeqA.sh” must run. This action will compute “AlcoR simulation” and generate a synthetic sequence using a uniform distribution through a sub-program that uses a linear congruential random number generation (LCG) [Kroese et al., 2013]. This sequence (sample.fasta) contains eleven segments of synthetic DNA where some of which are exact, inverted or mutated copies. An illustration of this sequence’s low-complexity coordinates (lilac color) is added to Figure 2-A) as the ground truth (Truth). The details of the composition of this synthetic sequence are described at the Supplementary Subsection 2.2.

**Figure 2.**
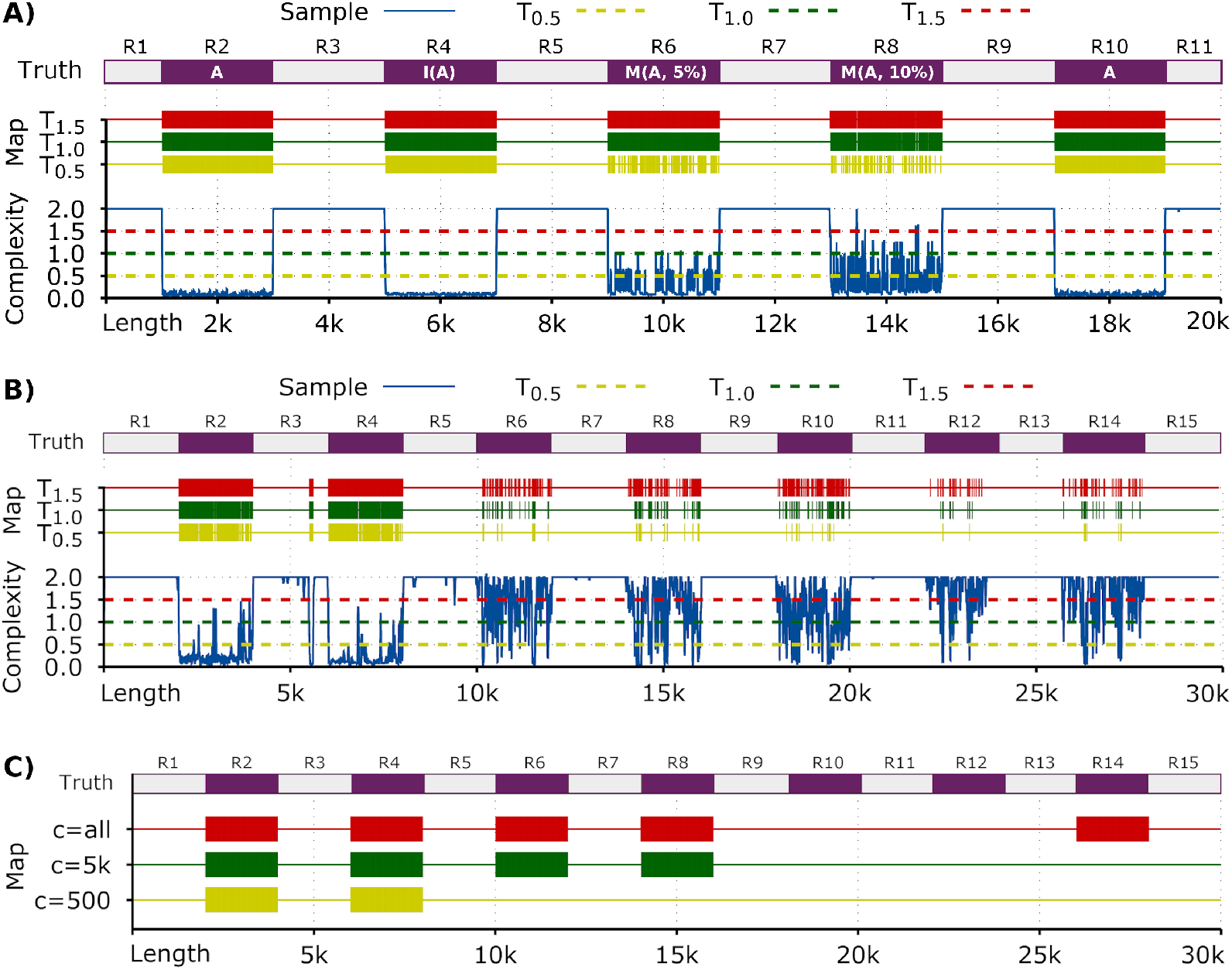
Low-complexity map and profiles for demonstrations A, B, and C. A) Minimal bi-directional complexity profile of a synthetic sequence (sampleA.fasta) and the corresponding map of the segmented LCRs below the respective threshold. In the Truth panel, the A region is used as a fixed random sequence for the following transformations: inversion (I) and mutation (M) with the respective percentage. The remaining regions are probabilistic random and unique. B) Minimal bi-directional complexity profile of the sampleB.fasta sequence and the corresponding map of the segmented LCRs below the respective threshold. In the Truth panel, the regions are according to the simulations and transformations described above. C) Low-complexity map of the sampleC.fasta sequence with the segmented regions below the 1.0 threshold for different *c* values (yellow - 500, green - 5,000, red - whole memory). In the Truth panel, the regions are according to the simulations and transformations described above. All the Truth panels have been added to the image.

This demonstration aims to benchmark the method, namely in the capability to find these regions, namely through similarity despite the transformations of inversion or mutation. For computing this demonstration, the “./DemoA.sh” script must run. This action will compute the “AlcoR mapper” for three thresholds (0.5, 1.0, and 1.5). The smoothing window is set to 10, and a context model of order-13, prepared to deal with inversions, is used in cooperation with a substitution-tolerant context model (allowing five substitutions in a context of 13) of the same order. Accordingly, the inner parameterization for each threshold is set to a model with 13:50:0:1:10:0.9/5:10:0.9, a window of 5, and a threshold of 1. Additional information on the parameters and meaning is available in Supplementary Section 3.

This tool will compute equation 4 and the respective segmented regions. Then, Gnuplot is used to generate the plots automatically from the script. The output plot is a PDF file with the name outA.pdf containing the minimal bi-directional complexity profile and the corresponding horizontal map of the segmented LCRs for each threshold.

As depicted in Figure 2-A), all the LCRs are accurately mapped for the thresholds *T*_1.5_ and *T*_1.0_, including the inverted region and both copied regions with substitutions. Identifying these regions is more challenging for the lower threshold, despite their good performance, assuming the high level of mutations that R6 and R8 contain.

#### 2.1.2 Demonstration B

Although the performance of Demonstration A seems to be quite convincing, the application in natural sequences has additional challenges. One of the challenges is the presence of a non-uniform nucleotide content. Therefore, instead of a maximum average complexity for encoding each base of log_2_(Θ), this value will be approximately at least equal to or less than the Shannon entropy [Shannon, 1948]. Therefore, the threshold needs to be approximated to half of this value. Usually, visual inspection and optimization after the first run provide efficient results, but this process can also be automatic.

Another challenge is the characteristics of non-stationary content of higher complexity regions, usually with the presence of peaks that have a mixed vertical direction (the same as in region R8 but of average higher complexity). This challenge requires an adaptive decrease of the smoothing parameter. However, this parameter solicits a trade-off. Lower the smoothing parameter, lower the precision of the regions below a given threshold, while higher the parameter, higher fragmentation and possibly lower accuracy in detecting the LCRs. Nevertheless, good precision and accuracy are achievable with a proper balance.

Accordingly, in this demonstration, a mixture of regions from synthetic and generated sequences is used recurring to context models trained with a non-synthetic viral sequence. For accessing the folder with the contents to perform this demonstration, the directory must be changed to “demoB”. The “GenSeqB.sh” is a script that will create a FASTA file (sampleB.fasta) with the conjunction of several synthetic and non-synthetic sequences. For generating this file, the “./GenSeqB.sh” command must run. This action will compute “AlcoR simulation” and generate the sequences containing fifteen segments of synthetic DNA that some of which are exact, inverted, or mutated copies.

An illustration of this sequence is provided in Figure 2-B as the ground truth (Truth). The details of the composition of this synthetic sequence are described at the Supplementary Subsection 2.3.

For computing this demonstration, the “./DemoB.sh” script must run. This action will compute “AlcoR mapper” similar as in Demonstration A but instead using the window size of 20 (the whole sample is larger). The output is the PDF file outB.pdf with the content provided in Figure 2-B.

This demonstration results include some examples with very high mutation rates, for example, a deletion mutation of a particular symbol with a probability of 0.15 and an additional mutation of a specific symbol with a probability of 0.1. Moreover, the sub-sequences contain substitution mutations, inversions, and origins from different sources. Despite the complexness of the challenge, all the LCRs have been identified. A small low complexity region has been additionally identified in the R3 mainly because this sequence belongs to the HHV6B, which contains a low-complexity sequence. Despite the correct identification, the tuning of the filter window size can improve the results, for example, by enlarging the mapped LCRs.

#### 2.1.3 Demonstration C

Demonstrations A and B showed the capability of the methodology to find LCRs with lower to higher mutation rates. However, a question remains: how to split local from distant LCRs? This demonstration answers this question by computing the method using different cache histories (*p*). Several sub-sequences with local and distant LCRs are simulated and modeled using natural sequences for this benchmark.

For accessing the folder with the contents to perform this demonstration, the directory must be changed to “demoC”. The “GenSeqC.sh” will create a FASTA file (sampleC.fasta) with the sequences while assuming different low-complexity levels and distances. For generating this file, the “./GenSeqC.sh” command must run. This action will compute “AlcoR simulation” and generate a sequence with the mentioned characteristics.

An illustration of this sequence is provided in Figure 2-C as the ground truth (truth). The details of the composition of this synthetic sequence are described at the Supplementary Subsection 2.4.

For computing this demonstration, the ‘./DemoC.sh” script must run. This action will compute “AlcoR mapper” similar as in Demonstration A and B but exclusively to map the LCRs for different *c* values. The output is the PDF file outC.pdf with the content provided in Figure 2-C.

This demonstration shows the potential of this application for mapping and visualizing LCRs with low (500 bases), medium (5,000 bases), and high distances (all bases) between patterns. Region R4 is a copy of R2, and a high redundancy content characterizes R2, thus enabling the low memory be enough to map these regions. Regions R8 is a copy of R6 with high statistical entropy, naturally only mapped by medium and large *c*. Finally, the R14 is also a copy of R8 and R6, enabling the detection for *c >* 10*k*, where *c* =all provides this capability. As in demonstrations A and B, this demonstration shows the automatic and accurate identification of the whole regions containing low-complexity but, in this case, using different distances between low-complexity patterns. For example, while local patterns are associated with regulatory regions, distant patterns can account for similar genes, or segmental duplications [Išerić et al., 2022, Mouakkad-Montoya et al., 2021].

As mentioned before, AlcoR can be applied to other types of sequences as long as it respects the FASTA format. To prove this, we provide the same demonstration for amino acid sequences, including the simulation, mapping, and visualization of the respective LCRs. This process can be replicated using the instruction at Supplementary Section 2.4.1. The results show the same output map despite the difference in the sequences used and the automatic calculation of the threshold using the process described in Equation 5.

### 2.2 Biological Data

For testing AlcoR exclusively in biological data (no synthetic data), we use the reference genomes from nine Human Herpesvirus (HHV), namely the Herpes-Simplex Virus 1 (HSV-1, NC 001806.2), Herpes-Simplex Virus 2 (HSV-2, NC 001798.2), Varicella-Zoster Virus (VZV, NC 001348.1), Epstein-Barr Virus (EBV, NC 009334.1), Human Cytomegalovirus (HCMV, NC 006273.2), Human Herpesvirus 6A (HHV6A, NC - 001664.4), Human Herpesvirus 6B (HHV6B, NC 000898.1), Human Herpesvirus 7 (HHV7, NC 001716.2), Kaposi’s sarcoma-associated herpesvirus (KSHV, NC 009333.1). Using these genome sequences, we compute the low-complexity maps and manually align them with the structural images retrieved from the NCBI sequence viewer tool (NCBIv) at the NCBI portal for comparison purposes [Rangwala et al., 2021, Camacho et al., 2009, Altschul et al., 1990]. The AlcoR computation map can be reproduced by running the script “RunHerpesvirus.sh” at the folder “herpesvirus”.

Figure 3 provides the low-complexity map of each Herpesvirus sequence included in this sample with the segmented regions below the 1.2 thresholds for local (red) and distant (green) regions and ignoring regions below 50 bases. Running the low-complexity maps with AlcoR, including the generation of the figure map for the whole Herpesviruses, took approximately 10 seconds using the computer characteristics described above. All the regions that the NCBIv mapped were reported by AlcoR (except for the final region in KSHV). For this final region, we changed the threshold for a lower value but did not have these regions considered as low-complexity. Also, we visually inspected this region and besides a tendency for a higher content of GC, we did not find any relevant low-complexity pattern. Moreover, we extracted the last 1000 symbols that contain this region highlighted by the NCBIv and measured the Normalized Compression (NC) [Pratas and Pinho, 2017] with GeCo3 [Silva et al., 2020] using custom optimized parameters (and ignoring the information header with the models) reaching a NC minimum of approximately 0.87 (repetitive values are close to zero and self-dissimilar values are close to one). This value indicates that there are no relevant low-complexity patterns in this subsequence. Potentially, this region may be underestimated, as an information quantity, by the NCBIv tool through the distribution of the nucleotide content.

**Figure 3.**
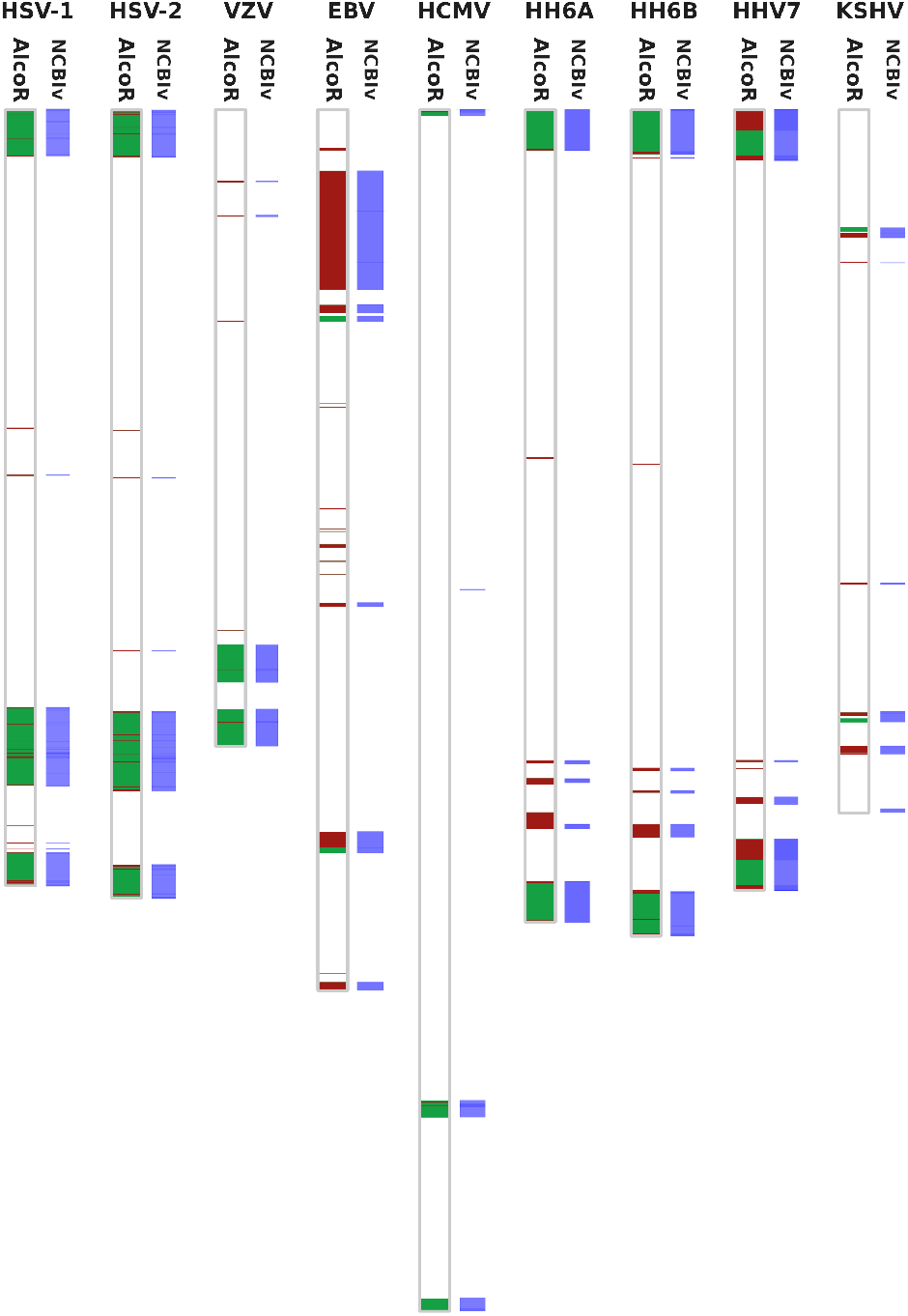
Low-complexity map of each Herpesvirus sequence included in this sample with the segmented regions below the 1.2 threshold for different *c* values (red - 5k symbols, green - whole memory). Red regions overlap green regions. In each viral sequence, the panel at the right describes the map produced by localizing repeating regions using the NCBI sequence viewer (NCBIv). The NCBIv panel has been added to the image. The maps are proportionally in scale according to the length of the sequences.

The remaining reported regions are consistently localized in identical coordinates, as shown in Figure 3 with the horizontal alignment between the maps. Additionally, AlcoR identified several more regions than the NCBIv tool. The main justification for these identified regions is given by the mixture of models and algorithmic methodologies that AlcoR contains, providing substantially higher sensitivity without compromising the efficiency of the computation regarding computational resources (memory and time) and, more importantly, without underestimation.

However, notice that the direct comparison of these tools is unfair. The parameters used in the NCBIv were the default (k-mer of 28 and threshold of 0.05), and by optimization, several of the regions that AlcoR mapped potentially will also be mapped by the NCBIv. However, the main reason for this unfair comparison is that these algorithms have different purposes, although they can be approximately compared for specific cases. The NCBIv tool can compute repetitive patterns through alignments using a single k-mer model; however, these are only a subgroup of LCRs. Additionally, the AlcoR tool can retrieve additional structural, distant, and algorithmic characteristics using multiple models with different capabilities. Nevertheless, we used the NCBIv tool specifically as a control.

Supplementary Table 1 provides information regarding the cumulative number of bases and respective sequence proportion considered by AlcoR as local and the overall (local and distant) LCRs. These results are the automatic computation of AlcoR that can be reproducible using Supplementary Section 2.5.

### 2.3 Large-scale Data

The computational and visualization of LCRs in large-scale data (*>* 100 MB) is more challenging because of the diversity of low-complexity patterns, the size of the sequences, and the required higher computational resources (computational time and RAM) to cope with the memory of distant low-complexity patterns. For this purpose, the window size must be adapted to a higher value (usually to 5,000), the compression models set with a high-order context model (at least 13), and if the number of segmented regions is too high for visualization purposes, ignoring some small regions (usually below 5,000) provides effective descriptive maps.

Taking into account these standard parameters, we run AlcoR unprecedentedly in two whole genomes, namely the recent complete human genome [Nurk et al., 2022] and the haplotype-resolved chromosome pairs of a heterozygous diploid African cassava cultivar (TME204) [Qi et al., 2022]. The low-complexity maps have been computed for both genomes using local low-complexity (using a memory of at most 5,000 symbols) and the distant (or global) low-complexity. Since the local low-complexity (red color) is depicted on top of the complete low-complexity in the visualization, it enables us to visualize the whole LCRs as exclusively distant (green color).

The computation of the whole maps took approximately 203 and 88 minutes for the human genome and the cassava diploid genomes, respectively, using the computational characteristics described above. These runs can be additionally parallelized, providing substantial computational time savings. The maximum peak RAM was approximately 1 GB for both genomes. To replicate the human run, the folder “human” contains the scripts “GetHuman.sh” and “RunHuman.sh” that must run in this order. To replicate the cassava run, the folder “cassava” contains the scripts “GunzipCassava.sh” and “RunCassava.sh” that also must run in this order.

The low-complexity map for the whole human chromosomes is depicted in Fig. 4. As depicted, most of the local LCRs overlap previous unsequenced regions characterized by high repetitive nature, for example, segmental duplications, GC regions, telomeric regions, among others [Išerić et al., 2022, Vollger et al., 2022]. Many of these regions are localized in the same coordinates addressed by segmental duplication analysis [Vollger et al., 2022].

**Figure 4.**
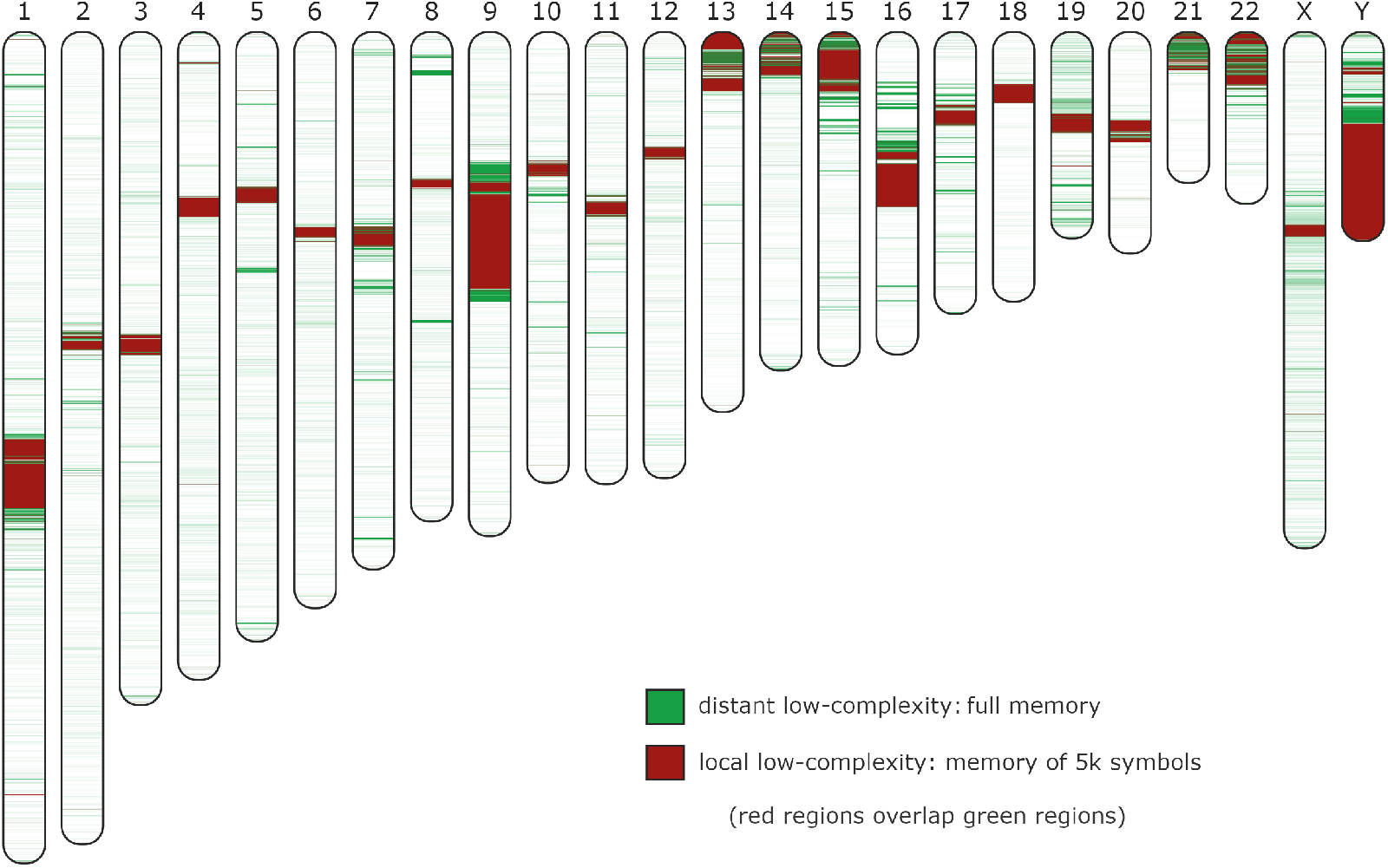
Human chromosomes low-complexity (redundancy) map using different memories. The green color stands for the identification of LCRs using a setup of a model that uses the full memory (cache history of all symbols), while the red color uses a memory of *p* = 5000 symbols. In the map, the regions with the red color overlap the green, therefore, regions marked with red are also green; the green regions that are visible stand for low-complexity related to distant patterns. Contiguous segmented regions below 5,000 symbols have been discarded from the visualization. All the chromosomes are in scale, with chromosome 1 (the largest) characterized by approximately 240 MB of length.

However, there are other regions that have been highlighted by AlcoR that are characterized by other sources, namely centromeric regions [Miga, 2020, Altemose et al., 2022]. Recently, a substantial structural diversity has been found in centromeric regions as human haplotype-specific evolution, while frequent SNPs remained conserved [Suzuki et al., 2020]. This finding shows the importance of having fast and sensitive tools to map these regions automatically for downstream analyses, especially in the new ongoing era of Telomere-to-telomere (T2T) assembly [Aganezov et al., 2022].

Supplementary Table 2 provides the overall low-complexity quantities for each chromosome according to its length for both local and distant patterns. The chromosomes that show the highest ratio of LCRs are: 1, 7, 9, 13, 14, 15, 16, 17, 19, 20, 21, 22, X, and Y. From these, the chromosomes with the highest levels of LCRs of local distant are: 9, 13, 15, 16, 19, 21, 22, and Y. On the other hand, chromosomes 2 and 6 contain the lowest ratio of LCRs.

Additionally, this method offers the characterization possibility of each region through the automatic extraction of each sequence by the respective coordinates. This means that each region can be extracted and analyzed or compared against well-known sequences, such as GC regions, telomeric, centromeric, TATA boxes, among others, and, after, the index color can be changed according to a desired value. This flexibility allows the creation of low-complexity maps with specific colours standing for different sequence characteristics.

The low-complexity map for the diploid cassava chromosomes is depicted in Fig. 5. It is interesting to notice that in several chromosomes, for example, in 1, 2, 4, 14, 15, 16, and 18, the number of low-complexity mapped regions is higher in one part of the chromosome than the other. This finding is also congruent in both the haplotypes. The questions that emerge are if these LCRs are smaller in a chromosome harm or some chromosomal harms contain more repetitive parts than the others and if this distribution is related to the organization of the genomes, rearrangement probability, and speciation.

**Figure 5.**
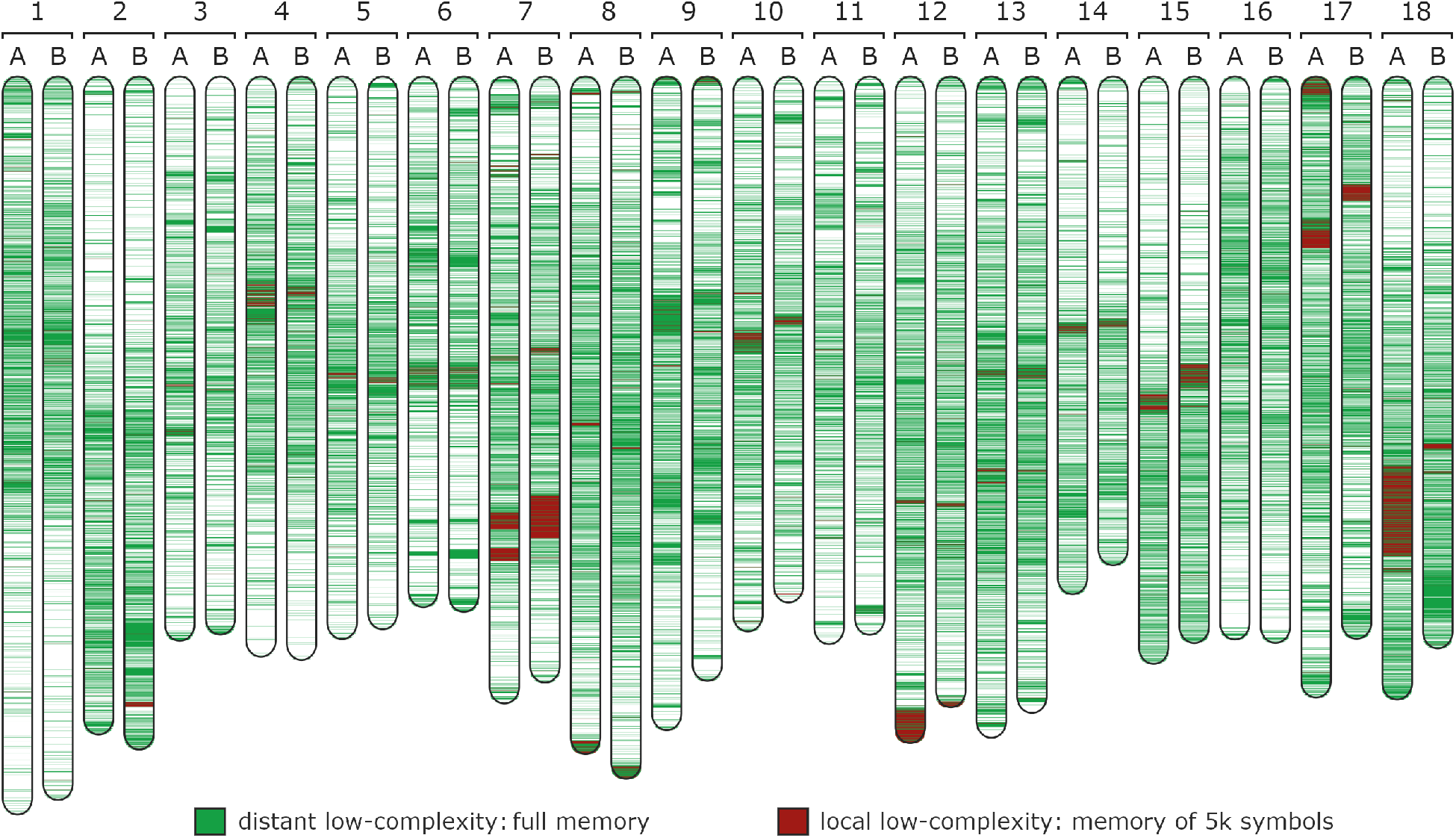
Diploid Cassava (TME204) chromosomes low-complexity (redundancy) map using different memories. Each chromosome number (1 to 18) contains the respective haplotype (A or B). We used 0.6 as the threshold because of the higher global low-complexity of this plant genome. The green color stands for the identification of LCRs using a setup of model that uses the full memory (cache history of all symbols), while the red color uses a memory of *p* = 5000 symbols. In the map, the regions with the red color overlap the green, therefore, regions marked with red are also green; the green regions that are visible stand for low-complexity related to distant patterns. Contiguous segmented regions below 5,000 symbols have been discarded from the visualization. All the chromosomes are in scale with chromosome 1 - haplotype A (the largest), characterized by approximately 44 MB of length. Although the labels have been manually added, the full map has been generated automatically using AlcoR.

In each pair of chromosomes, there are an extensive number of LCRs that, by a horizontal visual alignment between the haplotypes, depict an extensive number of visible vertical changes. Specifically, if the sequence pairs were very similar, the low-complexity map would be highly similar; this case shows many differences in all chromosome pairs. These changes are increasingly visible in the regions identified as local low-complexity (red regions). These findings are in accordance with the recent analysis of the extensive number of rearrangements found in this genome, namely through the comparison against previous references and between haplotypes [Qi et al., 2022].

Moreover, Cassava, like some other plant species, has “atypical” centromeres [Cuacos et al., 2015]. As in [Qi et al., 2022], we observed that in some chromosomes the number of LCRs was higher in one chromosome arm than the other, such as in extreme cases of chromosome 2 and 18, where the pericentromere and centromere region extends along one chromosome arm.

Supplementary Table 3 provides the overall low-complexity quantities for each pair of chromosomes according to their length for both local and distant patterns. We are able to notice extensive redundancy differences between the haplotypes. Specifically, the most obvious difference is in the local low-complexity percentage between haplotype A and B in chromosome 18. Supplementary Figure 1 enhances this difference that can only be marginally approximated in chromosomes 12 and 17.

We also notice two large local LCRs in the tips of haplotype A in chromosome 12 (end-tip) and 17 (beginning-tip). For example, the end-tip in chromosome 12 and haplotype A has been shown to be dissimilar relatively to reference AM560 v8.0 assembly and the haplotype B of TME204. The AlcoR method shows that this large segment is characterized by an approximate continuous local repetitive nature (below 5k symbols). Using AlcoR extract, we extracted this segment and noticed that it contains a large number of approximate copies of the sequence “TTTTGGGATGGAAGCTGTTAGTCCGAAATCGGGCACCGGATGTAACAATGATGATAGACTTGGCAG”. A BLASTn search (in 1-4-2023) of this sequence did not report any significant similarity, including the optimization with the increase of sensitivity, mismatch to 4/5, and the unfiltering of LCRs. This repetitive region is approximately 3% of this haplotype chromosome. An additional analysis (the comparative analysis that can be reproducible using Supplementary Subsection 2.6.2) on similar sequences in the remaining cassava sequences showed similarity with some regions of chromosome 17 (haplotype A) in nearly 1% of the chromosome (interestingly, not found in haplotype B), followed by chromosome 12 (haplotype B) with nearly 0.2%, and also in chromosomes 8 (haplotype A and B) with nearly 0.15% and 0.13%, respectively.

For benchmarking purposes, we now compare AlcoR against RepeatModeler and RepeatMasker [Flynn et al., 2020, Smit, AFA and Hubley, R and Green, P] in the task of sequence masking. Notice that this comparison is only indicative because the methods contain different specifications and, thus, the outcome can be substantially different.

From the AlcoR LCRs mapped regions, 68% and 70% of completely (100%) overlapped with those masked by RepeatModeler+RepeatMasker in A (Haplotype 1) and B (Haplotype 2), respectively. In both haplotypes, 6% AlcoR masked regions were not masked by RepeatModeler+RepeatMasker. The rest were partially overlapped.

The 6% AlcoR specific regions were likely caused by the parameter setting of RepeatMasker. These sequences were mostly very short, with low GC (Supplementary Figure 2). The longer sequences were simple tandem repeats. Since RepeatMasker was set with “-nolow -norna”, it is expected that simple tandem repeats and low complexity (polypurine, AT-rich) regions were not masked.

Among the regions masked by RepeatModeler+RepeatMasker, 3% in both haplotypes were not masked by AlcoR. These regions were masked by similarity to 1260/1256 repeat families predicted by RepeatModeler, of which only two repeat families (rnd-1 family-558 and rnd-1 family-598) were totally missed by AlcoR. The reason being is that AlcoR only mapped at a chromosome level for visualization purposes. Although this type of analysis can be performed by smash++ [Hosseini et al., 2020], AlcoR can also perform the whole genome analysis without affecting the processing time if the chromosomes are in a single FASTA file (for example, concatenated).

All in all, AlcoR is valuable in quickly masking LCRs in newly assembled genome sequences and visualising the LCRs. Specifically, AlcoR is highly-sensitive and can be used in comparative genomics for fast sequence complexity visualization. Moreover, AlcoR does not depend on software dependencies, unlike RepeatModeler. AlcoR is also time and resource-light, unlike the other tools. Moreover, it also contains additional tools for simulation, extraction, and to compute statistical information.

## 3 Discussion

Alignment-based methods usually offer an intuitive and enhanced local resolution that prevails in the comparative analysis of specific features. However, with the substantial increase in the development and availability of alignment-free methods [Vinga and Almeida, 2003, Reinert et al., 2009, Wan et al., 2010, Zielezinski et al., 2017, 2019], these alignment-free methods have evolved from unrolling an alternative way to provide clear advantages that would otherwise be complex to achieve using feasible computational resources, mainly in large-scale data. There are many examples, for instance, the SpaM [Morgenstern, 2021] and Prot-SpaM [Leimeister et al., 2019], that are providing the calculation of a higher data volume with similar or improved accuracy. Another example is the fast metagenomic composition characterization that can be achieved by these methodologies with high accuracy both for extant [Wood et al., 2019] and ancient [Pratas et al., 2018] samples.

Moreover, alignment-free methods based on lossless data compression offer a clear advantage: the absence of underestimation through the approximation of the Kolmogorov complexity. This property is essential to avoid false positives when the sensitivity is higher, especially when dealing with the identification of LCRs.

Therefore, the automatic identification of these LCRs without underestimation provides a sensitive and fast complementary approach to minimize the set of regions that have a higher probability of containing local errors and structural miss-assemblies in initial draft assemblies [Mc Cartney et al., 2022].

The recent development of the Telomere-to-Telomere (T2T) sequencing and assembly methodologies are providing the reconstruction of the full sequences, including the LCRs that previously were challenging to assemble because of the limited capability and assembly ambiguity that short reads provide [Treangen and Salzberg, 2012]. This technology is extremely valuable to provide completeness of large genomes and well as the ability to solve haplotype characterization. Ironically, this technology is solving the high-complexity task beyond the sequencing and assembly of LCRs, which are the main gaps that large sequence genomes, such as the human genome, previously contained.

In this article, we have proposed AlcoR, offering a convenient, fast, and easy way to automatically produce multiple low-complexity maps (and coordinates retrieval) of small-to-large sequence species that are being reconstructed with this technology without recurring to any additional tool. The AlcoR methodology is made available at the ideal time since the quality of these complete sequences is now satisfactory across the entire sequence, including in repetitive or redundant sub-sequences. From the cassava analysis in this article, we can verify a high variability of LCRs between chromosomal haplotypes that before were thought to be in a smaller quantity.

Another area where LCRs are currently being seen as fundamental is viral integration. This area addresses the potential capability of viruses to integrate the host genome, where some are particularly associated with tumorigenesis [Bodelon et al., 2016]. Therefore, viral integration is a probabilistic genetic marker for discovering virus-caused cancers and inferring cancer cell development [Cantalupo et al., 2018].

There are several computational tools to predict viral integration [Pischedda et al., 2021, Rajaby et al., 2021, Cameron et al., 2021, Chen et al., 2019]. However, only recently a methodology concerning LCRs (namely repeats) has been developed [Rajaby et al., 2021]. Specifically, inaccuracy is caused by the inconsistent alignment of reads in repetitive regions. Since integration sites vary according to the virus, host (usually in telomeric regions), and unknown factors, the availability of AlcoR to map and localize LCRs in both virus and host is significant; this allows the selection, validation, and knowledge building on this highly important thematic especially with the surprisingly high number of viral genomes residing in human tissues [Pyöriä et al., 2023].

The AlcoR package also contains a new FASTA simulation tool. The AlcoR mapper could not be adequately tested without the simulation tool because of its mapping singularity capability. The AlcoR flexible simulation tool allowed us to successfully validate the mapper by generating the data according to ground truth. Notice that validating AlcoR against other tools is unfair because the mapper is designed to dismiss false positives and enables finding LCRs containing different characteristics and specific potential algorithmic expressions. Nevertheless, both the AlcoR simulation and mapping tools have been validated in a complementary way. The AlcoR simulation tool would not be properly validated without a tool to detect LCRs. At the same time, the AlcoR mapper tool would also not be properly validated without the simulation tool. Therefore, this duality represented in this article was essential.

The final remark of low-complexity mapping is related to optimizing reference-free data compression tools. Identifying regions with similarity within a sequence enables clustering regions with the same characteristics, especially for distant LCRs. This means that AlcoR can be used to map distant LCRs followed by grouping and using a data compressor with lower RAM to increase the lossless compression of genomic or proteomic sequences. Naturally, this requires using side information to describe the regions and ensure a complete and lossless decompression. However, notice that the decompression side does not require the re-detection of these regions; hence, it provides a faster decompression process. Therefore, the development of tools such as AlcoR contributes to developing higher-ratio data compression tools while maintaining the same or smaller computational resources in the lossless decompression process.

## 4 Conclusion

This article proposed an alignment-free method for the simulation, mapping, and visualization of LCRs in FASTA data. The implementation of the method has been provided in an open-source tool (AlcoR) without any dependencies on external software. The AlcoR method uses parallel bi-directional complexity profiles. It is fully automatic, where the input consists of a FASTA file, and the output provided are coordinates and an SVG image map and file(s) with the respective coordinates. AlcoR can successfully mask sequences with high sensitivity and without underestimation.

Included in this tool is the additional capability to simulate FASTA sequences with different characteristics, namely using reference-based and reference-free approaches as well as the application of custom degrees of mutations. The AlcoR mapper can distinguish between local and distant LCRs without underestimation and is optimized and efficient to be applied in small- to large-scale sequences.

The tool has been successfully benchmarked using synthetic, nearly synthetic, and natural sequences, providing whole LCRs identification using efficient computational resources. We applied AlcoR in large-scale data, unprecedentedly providing a whole-genome low-complexity map of a recent complete human genome and haplotype-resolved chromosome pairs of a heterozygous diploid African cassava cultivar. Additionally, we extended the analysis to identify some insights into repetitive cassava-specific sequences.

With the current Telomere-to-telomere (T2T) sequencing and assembly methodologies, AlcoR offers a convenient, fast, and easy way to automatically produce multiple low-complexity maps of species that are being sequenced and assembled with this technology without underestimation and recurring to any additional tool.

## Acknowledgements

The authors thank the Finnish Computing Competence Infrastructure (FCCI) for supporting this project with computational and data storage resources.

